# Resource scarcity increases foraging activity despite thermal risk in an arid-adapted bird

**DOI:** 10.1101/2025.11.13.688088

**Authors:** Charlotte Christensen, Kennedy Sikenykeny, Damien R. Farine

**Affiliations:** University of Zurich, Department of Evolutionary Biology and Environmental Studies, Winterthurerstrasse 190, 8057 Zurich, Switzerland; Mpala Research Centre, P.O. Box 555, Nanyuki 10400, Kenya; Max Planck Institute for Animal Behavior, Am Obsterberg, Radolfzell 78315, Germany; Nairobi University, Faculty of Veterinary Medicine, Department of Clinical Studies, P.O. Box 29053-00625, Kangemi, Loresho-Ridge Road off Kapenguria Road, Nairobi, Kenya; Australian National University, Research School of Biology, 134 Linnaeus Way, Acton ACT 2601, Australia

**Keywords:** accelerometers, arid systems, birds, drought, foraging ecology, resource scarcity, sublethal costs, thermal homeostasis vs. foraging trade-offs

## Abstract

Behavioural flexibility is considered key for species to cope with the effects of climate change. Periods of resource scarcity, such as during climate change-driven droughts, may force animals to increase their foraging activity to meet energetic demands. However, doing so may increase the thermal risk inherent to the trade-off between foraging and exposure to higher temperatures. We use biologging data (internal body temperature and activity budgets classified via accelerometers) from a population of wild vulturine guineafowl (*Acryllium vulturinum*) to quantify this trade-off across varying resource availability (quantified through NDVI) over 2.5 years, including a period of severe drought. We start by linking internal body temperature measurements to external operative temperature to identify a ‘thermal threshold’—where body temperature reaches a maximum—at an external temperature of 33.8°C. We interpret this threshold as the point where birds must prioritise heat dissipation (physiologically and/or behaviourally) to avoid hyperthermia. Relevantly, we find that a larger part of the day exceeds this external ‘thermal threshold’ as resources decline. We then show that guineafowl feeding activity positively correlates with resource scarcity, suggesting the birds compensate for diminishing resources by increasing foraging time investment, despite thermal risks. However, the compensatory foraging does not appear to buffer the birds against food shortfalls, as birds still lose weight as resource scarcity increases. Sublethal effects (e.g., body mass loss) of compounding environmental stressors have fitness consequences (e.g., postponement of breeding, as was noted in our population). Our results could point to an evolutionary mismatch; the projected intensification of resource scarcity and higher temperatures under climate change may alter trade-offs in such a way that existing physiological and behavioural responses to cyclical dry seasons are no longer adaptive, in vulturine guineafowl, and in other arid-system organisms more broadly.

## Introduction

Climate change is set to affect food availability, with extreme and severe weather events bringing about resource bottlenecks (Maron *et al*. 2015). Droughts in particular pose a challenge in this regard, and are projected to intensify in duration and intensity, especially in arid and semi-arid regions (Cook, Mankin & Anchukaitis 2018; Haile *et al*. 2020). Not only do droughts reduce food abundance (Dickman *et al*. 2011), they also cause food patches to become further dispersed from one another (Macdonald & Johnson 2015) and remaining food to be less nutritious (Lambert & Rothman 2015) or harder to obtain (Gesquiere *et al*. 2008). Drought-driven resource scarcity is further exacerbated by rising temperatures (Mukherjee, Mishra & Trenberth 2018), creating a landscape that is increasingly challenging even for arid-adapted wildlife (du Plessis *et al*. 2012; Buchholz *et al*. 2019; Funghi *et al*. 2019; Riddell *et al*. 2019; Fuller *et al*. 2021; Tatler *et al*. 2021). A first line of defence for dealing with harsh environmental conditions—such as those experienced during droughts—is behaviour (Buchholz *et al*. 2019; Wong & Candolin 2024), because unlike morphological change, behaviour can be changed moment-to-moment to navigate a dynamic resource and thermal landscape. Nevertheless, behaviour can only be expressed within certain physiologically permissible boundaries (e.g., thermal and dehydration tolerance: Fuller *et al*. 2016; Funghi *et al*. 2019) and with every behavioural decision comes a trade-off.

Foraging is a central activity in many animal’s time budget; failing to meet nutritional requirements leads to starvation and eventually death (Dunbar, Korstjens & Lehmann 2009). How should animals allocate their time to maximize their energetic intake (necessary to maintain body-condition) while minimizing immediate risk, time and energetic investment (Abrams 1991)? As resources become scarcer, many animals devote more time to foraging. For example, during dry seasons, greater kudu (*Tragelaphus strepsiceros*) (Owen-Smith 1994), yellow baboons (*Papio cynocephalus*) (Gesquiere *et al*. 2008) and chimpanzees (*Pan troglodytes*) (Murray, Eberly & Pusey 2006) have been noted to diversify or shift their diet and spend more time foraging. In marine systems periods with low fish and krill are associated with similar upticks in foraging effort in African penguins (*Spheniscus demersus*) (Campbell *et al*. 2019) and marbled murrelets (*Brachyramphus marmoratus*) (Ronconi & Burger 2008). However, foraging activity, particularly under inclement conditions, carries physiological (Costa, Croxall & Duck 1989; Gesquiere *et al*. 2008; Kokubun *et al*. 2018; Van de Ven, McKechnie & Cunningham 2019) and opportunity costs (Cunningham, Gardner & Martin 2021), as well as risks. In the most extreme cases, investing in foraging can result in death by predation (Werner & Anholt 1993) or exhaustion of body reserves (Watter *et al*. 2019). Therefore, immediate self-maintenance must sometimes take priority over foraging (*sensu* ‘enforced resting time’: Dunbar, Korstjens & Lehmann 2009). This can include behavioural thermoregulation strategies in response to elevated temperatures (Hafez 1964) like seeking shade, pausing heat-generating activity (e.g., foraging, locomoting), or through evaporative heat loss (e.g., gular fluttering, feather raising in birds: Krishnan et al., 2023; exposing tongue in mammals: Campos & Fedigan 2009). During periods of resource shortage and higher temperatures, foraging activity therefore becomes diametrically opposed to maintaining safe body temperatures.

Capturing the dynamics of activity budgets in response to environmental change gives insights into an animal’s ability to cope – and ultimately a population’s viability – under climate change (Sih 2024). For example, we can monitor the increase in foraging investment to infer when foraging may be hitting a ‘hard ceiling’, whereby physiological limits do not allow for further increments. Baboons increased foraging up to a point during periods of resource scarcity, before plateauing and ‘switching strategies’ (e.g., started to rest during the hottest temperatures of the day: Hill 2006; Gesquiere *et al*. 2008). Similar upper thermal thresholds for foraging have been identified in other endotherms (Ostrowski, Williams & Ismael 2003; Tieleman & Williams 2002) and in ectotherms (Putman & Clark 2017; Roeder, Paraskevopoulos & Roeder 2022). If climate change-driven warming and resource scarcity increases as projected (Maron *et al*. 2015; Mukherjee, Mishra & Trenberth 2018), hitting these upper thermal limits is likely to happen more frequently and for more extended periods of time in the future. While higher temperatures can drive animals to avoid foraging during certain parts of the day (Levy *et al*. 2016), long-term resource scarcity may impose constraints that drive animals to abandon the ‘safe’ strategy that minimises heat stress, to instead prioritize meeting energetic demands (du Plessis *et al*. 2012). Indeed, as the climate warms, the viable time window for foraging activity may be narrowing to such an extent that sheltering from the heat is not an option (Fuller *et al*. 2016), particularly for those animals that are restricted to being active during the day (Hut *et al*. 2012) and/or are limited in when they can forage by predation pressure (Veldhuis *et al*. 2020; Vermeulen *et al*. 2024). In this scenario, monitoring foraging budgets alone may only reveal part of the energy intake *vs*. thermal homeostasis trade-off, as animals might be able continuing to forage but be incurring physiological costs. Generally, the ability to forage at sub-optimal temperatures (or in sub-optimal habitats) is seen as a ‘positive’ in terms of animals keeping pace with climate change through behavioural flexibility (Beever *et al*. 2017), but physiological data is needed to resolve whether there may be hidden, sub-lethal costs.

Here, we show how a diurnal, arid-adapted bird—the vulturine guineafowl (*Acryllium vulturinum*)–responds to decreasing resource availability (characteristic of dry seasons with low vegetation and insect abundance) as the trade-off between foraging activity and thermal homeostasis intensifies. Vulturine guineafowl are endemic to East Africa (BirdLife, 2024), where droughts present an increasingly pressing challenge (Haile *et al*. 2020). Previous studies have found that vulturine guineafowl expand their home-ranges during dry seasons (Papageorgiou *et al*. 2021), presumably in search for food. However, how foraging activity itself is modulated has not yet been tested. We first establish a ‘thermal threshold’ by combining data from internally implanted temperature loggers and external environmental loggers capturing operative temperature (the temperature as experienced by the organism: Dzialowski 2005). Endotherms, such as birds, generally maintain a constant body-temperature—within a relatively narrow range (Boyles *et al*. 2011)—but can show increases in body-temperature as external operative temperatures increase (Lovegrove, Heldmaier & Ruf 1991; Angilletta *et al*. 2010). As operative temperatures increase, we expect to see a point (thermal threshold) above which the birds start paying a disproportionately larger cost to maintain thermal homeostasis (Boyles *et al*. 2011; Rezende & Bacigalupe 2015). We then contextualise this thermal threshold by estimating what proportion of the day exceeds this threshold as resources decline. Finally, we test whether the increased foraging effort expressed by the birds buffers them against food shortfall, by investigating whether birds lose weight during resource scarcity. Based on long-term trends and future projections for East Africa, animals will face more periods of reduced food availability that will coincide with higher temperatures (Gebrechorkos, Hülsmann & Bernhofer 2019; Haile *et al*. 2020), raising questions about whether the behavioural responses expressed by animals who evolved in these harsh environments (and similar environments globally) are still adaptive (Sih, 2024).

## Methods

### Study system

We use data from a population of vulturine guineafowl that are part of a long-term project, the Vulturine Guineafowl Research Programme, at the Mpala Research Centre, Laikipia, Kenya (00.29°N, 36.90°E). Birds are fitted with solar-powered tags (15 g Bird Solar, e-obs Digital Telemetry, Grünwald, Germany, for details see: Papageorgiou *et al*. 2021) with an onboard tri-axial accelerometer (ACC). While we typically collect GPS data from >100 individuals at any given time, download limitations (He *et al*. 2022) only allow us to collect continuous ACC data from a subset of the population. Between 2021 and 2024, we collected ACC data for n=49 vulturine guineafowl (n=14 females, n=35 males) belonging to 17 social groups over a period of 644 days (between 2021-08-29 to 2024-03-08). Tags collected data for variable lengths of times (median: 171 days, range: 9-344 days; see Fig. S1 for monitoring time per bird), with 42/49 birds collecting data for > 1 month and 23/49 birds collecting data for > 6 months. Some birds experienced little variation in ecological conditions on account of the extended drought conditions between 2021-2023 (UNDRR, 2024; Fig. S1).

### Data collection

#### Accelerometer (ACC) data

To strike a balance between data collection, battery life, and download limitations, tags were programmed to collect ACC data between 06:20 and 11:30 local East African Time (EAT), which covers the morning foraging hours. ACC data were used to train a supervised machine-learning model to classify behaviours into six behavioural states. Details on behavioural classification methods have been previously described and biologically validated (Christensen *et al*. 2024). Here, we focus on ‘feeding activity’ (when birds actively peck at the ground or head-level vegetation) to quantify the amount of time the bird is extracting food. To calculate activity budgets from the second-by-second ACC data, daily feeding time was divided by total time with ACC data in the morning to obtain a feeding rate (Altmann 1974). Details on sampling regime can be found in *Section S1* of the *Supporting information*)*. Internal temperature loggers* As part of a previous project (Brandl *et al*. 2025), seven birds (not the same birds used in the accelerometer-analysis) were implanted with electrocardiogram (ECG) loggers that recorded heart rate and body core temperature for 6 seconds every 20 seconds between 02-03-2021 and 30-04-2022, from 05:00 to 20:00 EAT. From initial inspection of the data, one bird was identified as having outlier temperature readings relative to the others (see *Section S2* in the *Supporting information* for details). Our resulting sample size was therefore six birds for a total of 630 bird days (median: 108, range: 29-155 days of logging per bird).

#### Environmental data

We use Normalized Difference Vegetation Index (NDVI) data (resolution: 10 by 10 meter cells) calculated from European Space Agency Copernicus Sentinel-2 satellite images to assess change in vegetation quality (Pettorelli *et al*. 2005). NDVI values ranged from 0.15 to 0.52 in the study period (29-08-2021 to 08-03-2024; for details see *Section S3* in the *Supporting information*). NDVI correlates closely with vegetation quality (Ali *et al*. 2013) and insect abundance (Fernández-Tizón *et al*. 2020; Traba *et al*. 2022), including in savanna ecosystems (Pringle *et al*. 2007) such as the one we work in. As the guineafowl forage predominantly on grasses, seeds and insects (Del Hoyo *et al*. 1992; unpublished data KS thesis), we consider NDVI a good proxy for resource availability. To aid interpretability of the findings, we calculated resource scarcity as:

resource scarcity = 1 – NDVI

such that higher values correspond to greater resource scarcity and lower values correspond to greater resource availability.

#### Temperature data

We used temperature data from ten DS1922L iButton (Maxim Integrated) loggers (which take a temperature reading every 5 minutes) fitted into model guineafowl (hollow black iron cylinders with white dots) distributed across the home-range of the study population to estimate operative temperature (Dzialowski 2005). Model guineafowl were placed in pairs under cover and in the open to obtain data on temperature differentials (see Rozen-Rechels *et al*. 2025 for details), but here we used temperature measures from open areas as guineafowl predominantly use highly productive (open) glades for foraging (Young, Patridge & Macrae 1995). For the internal body temperature analysis, we took hourly means of the external operative temperatures (across the ten iButtons) and matched those to mean internal body temperature of each individual guineafowl on the same hour of the same day.

### Data analysis

All statistical analysis were conducted in R (version 4.1.1) and R studio (version 2024.04.2+764). The following packages were used for data structuring (dplyr, tidyr, lubridate), fitting generalized and linear mixed models (lme4, lmerTest, scales), and plotting (ggplot2, cowplot). Where linear mixed models were used, normality assumptions were verified visually using Q-Q plots of the residuals.

#### Relationship between internal body temperature and external operative temperature

Unlike heat stress experiments run in captivity where exposure to temperature, water, space and shade is controlled (e.g., Hamrita & Conway 2017), our data comes from a wild system in which the animals are moving freely and engaging in all activities, including heat-dissipating activities. While birds (and other endotherms) can maintain thermal homeostasis over large external temperature ranges, body temperature can increase with operative temperature (Lovegrove, Heldmaier & Ruf 1991; Angilletta *et al*. 2010). We hypothesized that the relationship may peak at some maximum operative temperature, with the maximum marking the point where the individuals must invest in thermoregulatory strategies to avoid risks of hyperthermia (Boyles *et al*. 2011; Rezende & Bacigalupe 2015). We note here that we did not make any assumptions about the thermoregulatory strategy; the mechanism could be physiological, behavioural or both (Angilletta *et al*. 2010). For details on how the ‘thermal threshold’ was calculated see *Supporting information, Section S4*.

#### Relationship between resource scarcity and feeding activity

To estimate the effect of resource availability on feeding activity, we fit a generalized linear mixed model with total feeding time relative to the total morning time as the response variable with a binomial error distribution and resource scarcity as the predictor. We fit sex as an interactive term with resource scarcity, as sex differences in feeding activity are plausible based on differences in body size (e.g., Ishikawa & Watanuki 2002; Lewis *et al*. 2005) and life history strategies (e.g., Powolny *et al*. 2014; Van de Ven *et al*. 2020). We fit bird ID as a random slope and date as a random intercept. We used this model to predict feeding activity values for each sex at incremental resource scarcity intervals (Fig. 2a).

**Figure 1.**
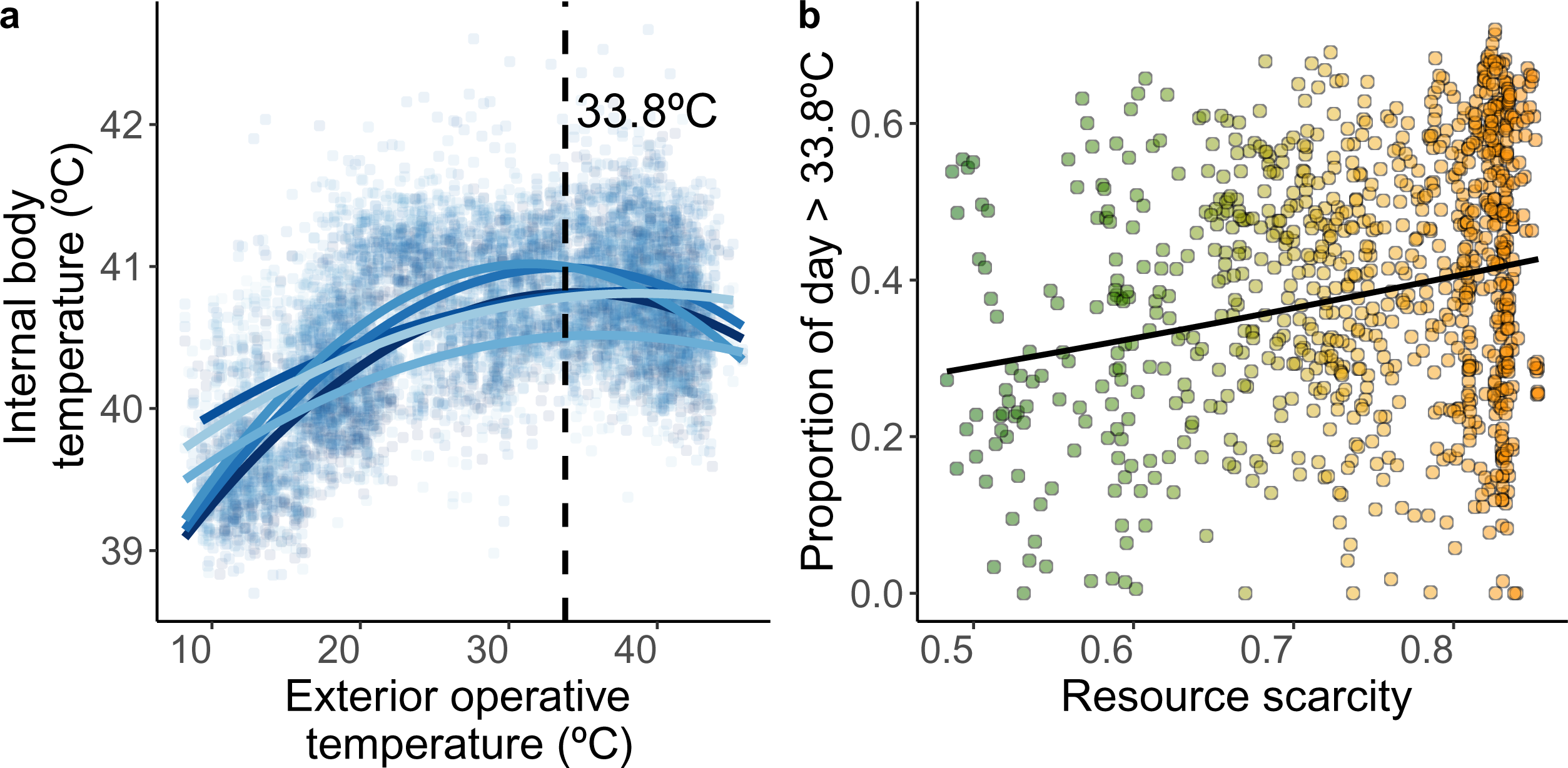
Birds must engage in behavioural or physiological strategies to maintain thermal homeostasis at high operative temperatures, and periods of resource scarcity correspond with a larger proportion of the day exceeding the critical temperature at which these strategies are necessary. (**a**) Relationship between hourly external operative temperature and internal body temperature (both in °C) from six vulturine guineafowl fitted with internal temperature loggers. Points are raw data points; blue lines are the fitted quadratic terms with each bird represented in a different shade. The dashed vertical line marks the maximum point of the curves, which the model revealed to occur at 33.8°C. This point indicates the operative temperature above which internal body temperature does not increase any further, meaning that birds are likely to engage in (physiological and/or behavioural) activities to maintain thermal homeostasis. (**b**) The proportion of the day (06:00 to 19:00) that exceeds the thermal threshold of 33.8°C increases with resource scarcity. Data are shown for the 2.5-year period where ACC birds collected data. Each point represents one day, and the colours represent increasing resource scarcity (from green to orange).

**Figure 2.**
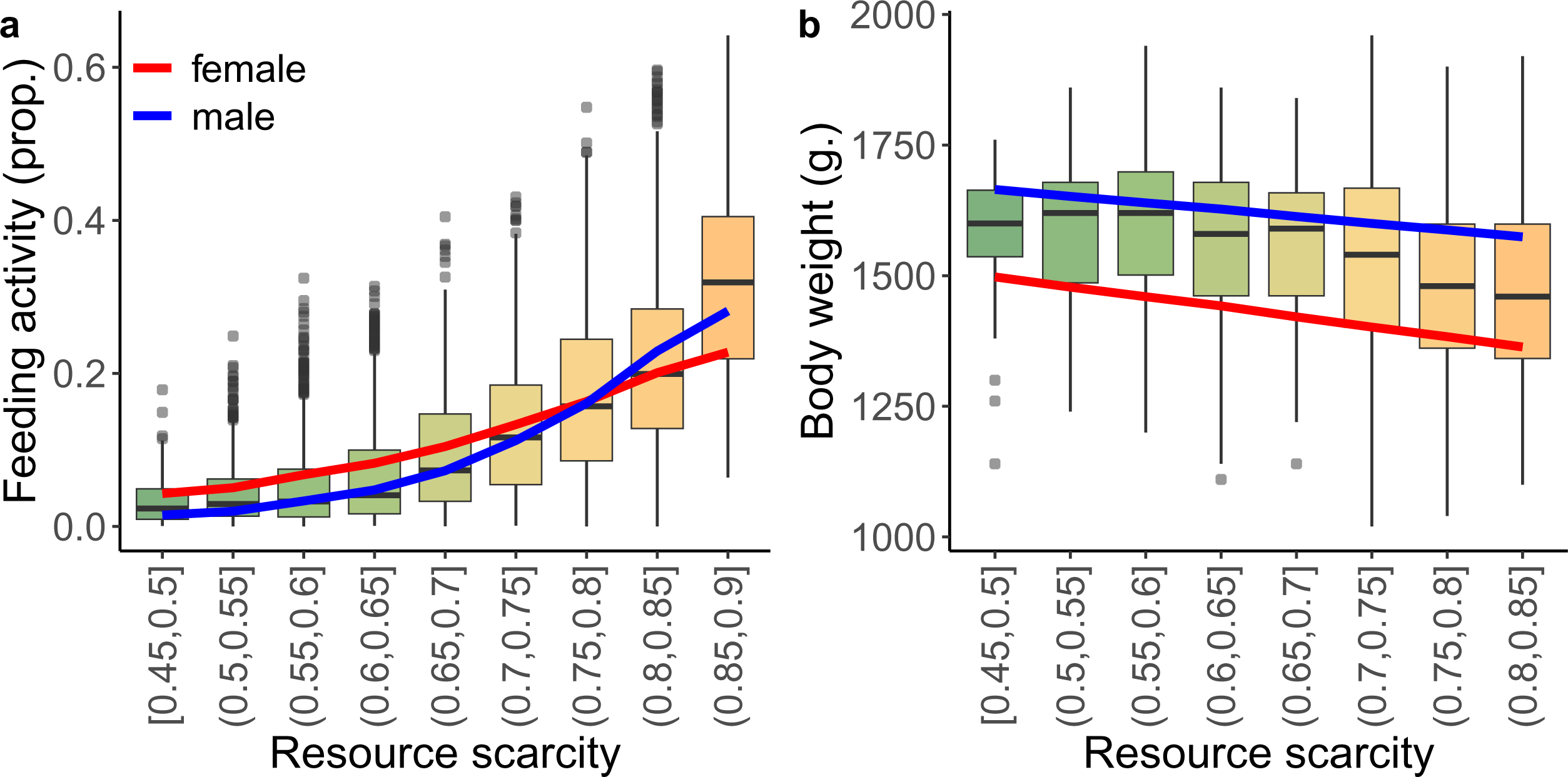
Birds substantially increase foraging effort when resources are scarcer, causing increasing thermal maintenance vs. foraging trade-offs that potentially drive a loss of body weight. (**a**) Daily morning feeding activity proportions for n=49 adult vulturine guineafowl over n=644 unique dates (n=7632 bird days). Boxplots show the raw data (binned in 0.5 increments for visualisation) while the lines represent male (blue) and female (red) feeding activity as predicted by the generalized linear mixed model in relation to resource scarcity. (**b**) Body weights from n=1948 measurements across n=1024 adult vulturine guineafowl in relation to resource scarcity. Boxplots show raw data (binned in 0.5 increments for visualisation) while lines represent male (blue) and female (red) feeding activity as predicted by the generalized linear mixed model in relation to resource scarcity. In both (**a**) and (**b**) the colour scheme from green to orange represents increasing resource scarcity (1-NDVI). In (**a**) resource scarcity corresponds to the same day as the feeding activity data, whereas in (**b**) resource scarcity is the average resource scarcity in the two weeks leading up to captures (when body weight was measured).

#### Relationship between resource scarcity and external operative temperature

To test whether resource scarcity coincided with higher external operative temperatures during the study period (n=907 days), we calculated the mean daily temperatures from the external temperature loggers over the daylight hours 06:00 and 19:00, when the birds are generally active (Klarevas-Irby & Farine 2024). We then calculated what proportion of the day exceeded the identified ‘thermal threshold’ for the birds of 33.8°C (see “*Internal body temperature peaks at a threshold external temperature*”). We fit two models with resource scarcity as the predictor: one linear model with mean daily external temperature as the response variable and one generalized linear model with proportion of the day >33.8°C as the binomial response variable.

#### Relationship between resource scarcity and body weight

To test whether resource scarcity correlates with body weight loss, we used a long-term dataset (2018-02-21 to 2023-11-15) of n=1948 weights from 1024 adult vulturine guineafowl (n=543 males and n=481 females) collected during captures. While, ideally, we would match feeding activity directly to weight loss/gain, this is not possible in our study system as birds need to be captured to be weighed (which only happens during routine capture 0–2 times per group per year, to fit colour bands and check tag and GPS/ACC-backpack condition). We fit a linear mixed model with weight as the response variable and resource scarcity as a predictor variable while controlling for bird ID, trapping location, and date as random effects. We fit sex as an interactive term with resource scarcity to account for consequences of different changes in foraging activity (Table 3) or physiological sex-differences in response to food intake/deprivation (e.g., Reda *et al*. 2024). Between 2022-06-03 and 2022-07-12 a high calorie feeding experiment took place (Ogino 2025) that affected the weight of the birds, hence data during this period and the following 6 months were excluded.

**Table 1.** Birds have a temperature threshold above which they likely use behavioural or physiological strategies to maintain thermal homeostasis. Linear mixed model outputs demonstrating the polynomial relationship between hourly external operative temperature and matched hourly internal body temperature between 05:00 and 20:00 local time of n=6 vulturine guineafowl fitted with internal temperature loggers across n=630 bird days (n=206 unique dates; n=7719 unique hours). Sex was fitted as fixed interactive effect with external operative temperature. Date and bird ID were fitted as random intercepts in the model. Significant terms are in bold.

|  | **Estimate** | **SE** | **t-value** | **P-value** |
| --- | --- | --- | --- | --- |
| **Intercept** | 39.237 | 0.067 | 583.663 | **<0.001** |
| **Poly(external temp.)1** | 4.749 | 0.108 | 43.763 | **<0.001** |
| **Poly(external temp.)2** | -3.369 | 0.105 | -32.064 | **<0.001** |
| **Sex(male)** | 0.135 | 0.095 | 1.412 | 0.197 |
| **Poly(external temp.)1\*Sex(male)** | -0.19 | 0.162 | -1.178 | 0.239 |
| **Poly(external temp.)2\*Sex(male)** | -0.138 | 0.156 | -0.885 | 0.376 |

**Table 2.** Periods of resource scarcity correspond to higher external operative temperatures and a greater proportion of the day exceeding a thermal threshold during which birds need to use strategies to maintain thermal homeostasis. Effect of resource scarcity on mean daily temperature (linear mixed model) and proportion of the day exceeding the thermal threshold of 33.8°C (generalized mixed model with a binomial distribution) across n=903 days. Significant terms are in bold.

|  | **Estimate** | **SE** | **t-value** | **P-value** |
| --- | --- | --- | --- | --- |
| *Mean daily temperature* |  |  |  |  |
| **Intercept** | 25.225 | 0.728 | 34.650 | **<0.001** |
| **Resource scarcity** | 6.811 | 0.972 | 7.006 | **<0.001** |
| *Proportion of the day exceeding thermal threshold (>33.8°C)* |  |  |  |  |
| **Intercept** | -1.757 | 0.013 | -132.17 | **<0.001** |
| **Resource scarcity** | 1.714 | 0.018 | 96.97 | **<0.001** |

**Table 3.** Birds increase feeding activity when resources are scarce. Generalized linear mixed model results showing the effect of resource scarcity on the morning feeding activity of n=49 adult vulturine guineafowl (from n=7632 bird days spanning 644 unique dates), with males experiencing a significantly greater increase in feeding activity in response to resource scarcity. The model controlled for date as a random intercept and bird ID as a random slope. Significant terms are in bold.

|  | **Estimate** | **SE** | **z-value** | **P-value** |
| --- | --- | --- | --- | --- |
| **Intercept** | -5.672 | 0.524 | -10.831 | **<0.001** |
| **Resource scarcity** | 5.225 | 0.688 | 7.598 | **<0.001** |
| **Sex(male)** | -2.975 | 0.665 | -4.472 | **<0.001** |
| **Sex(male)\*Resource scarcity** | 3.827 | 0.894 | 4.281 | **<0.001** |

## Results

### Internal body temperature peaks at a threshold external temperature

Fitting external operative temperature as a polynomial term significantly improved the fit of the model relative to including it as a linear term (X^2^= 1589.1, df=1, *p*<0.001, see *Supporting information, Section S4* for model building details and Table 1 for full model output), suggesting birds engage in physiological or behavioural strategies to maintain thermal homeostasis when the external temperature reaches a certain threshold. The calculated maximum body temperature across the six curves—where these compensatory mechanisms are likely to be engaged—corresponds to an external temperature of 33.8°C (Fig. 1a) and a mean internal temperature of 40.9°C.

### Periods of resource scarcity have higher temperature and a greater proportion of days exceeding the thermal threshold

We found a significant positive relationship between resource scarcity and mean daily temperature (Table 2). Over the course of the study period, the proportion of the day exceeding the thermal threshold of 33.8°C was also significantly higher during periods of resource scarcity, increasing by over 40% between the days with the least resource scarcity and the most resource scarcity (Table 2; Fig. 1b).

### Birds forage more when resources are scarcer

We found a strong positive effect of resource scarcity on morning feeding rates (Table 3, Fig. 2a). Over the course of the study period, we found an average increase of 17% in feeding rates during the periods of the highest resource scarcity relative to the lowest.

### Birds lose body condition when resources are scarcer

We found a negative effect of resource scarcity on body weights, with females losing more weight as scarcity increased (Table 4; Fig. 2b). Specifically, over the period spanning from lowest resource scarcity to greatest resource scarcity, males lost on average 90 g (or 5% of their body weight) while females lost on average 130 g (or 9% of their body weight). Males (mean±SD = 1621±121 g) were also significantly heavier than females (mean±SD = 1398±134 g) overall (Table 4).

**Table 4.**
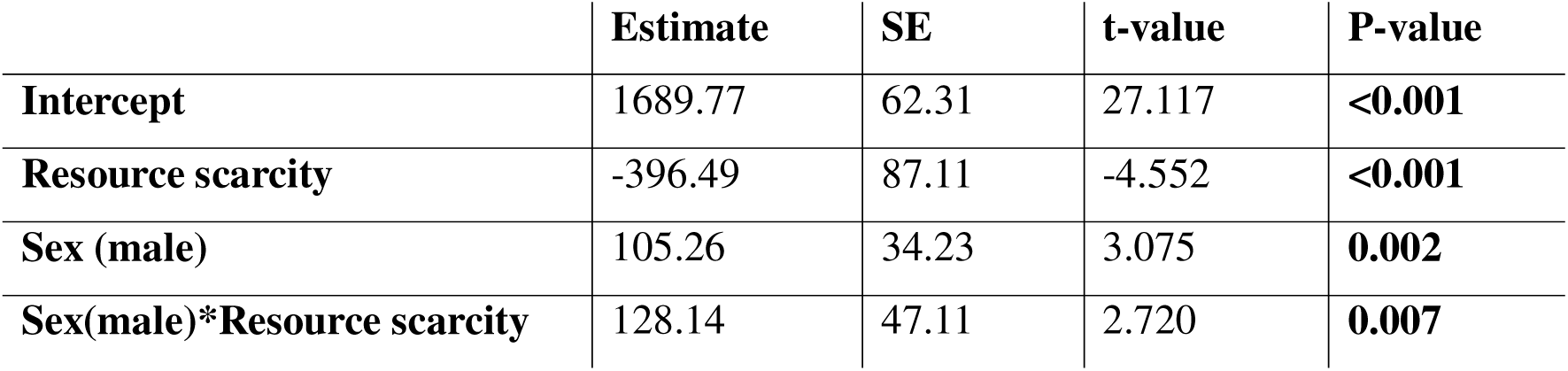
Birds lose weight when resource are scarce. Linear mixed model results showing the effect of sex and average resource scarcity across two weeks prior to measurement on the body weight of n=543 adult male and n=481 adult female vulturine guineafowl (n=1948 weights; n=95 unique dates), while controlling for date, trapping location and bird ID. Significant terms are in bold.

## Discussion

We show that vulturine guineafowl, an arid-adapted bird species, increase feeding activity as resources become scarcer. We also find that these harsher periods correspond to an increase in thermal risk. Specifically, we see a steep increase in the proportion of the day exceeding the thermal threshold (> 33.8°C) above which birds must invest (behaviourally or physiologically) in thermoregulation to maintain thermal homeostasis. As such, the birds face an increasingly challenge due to the foraging *vs.* thermal homeostasis trade-off; in the same period that the birds ramp up feeding activity (by c. 17%), they also face a shrinking suitable thermal window to forage in (>40% of the day exceeds 33.8°C). Our long-term data confirm that this trade-off has costs by showing that birds lose body-mass during periods of greater resource scarcity. Together, these results suggest that compensatory foraging does not appear to buffer the compounding challenges of reduced food availability and thermal risk as the birds incur sub-lethal costs. Finally, we found that whereas results were qualitatively consistent for males and females, males ramped up feeding activity more steeply and lost less weight relative to females as resources became scarcer.

When faced with greater resource scarcity, animals often increase their foraging effort. This investment can result in immediate rewards. For example, penguins that performed more foraging dives gained more body mass (Lescroël *et al*. 2021). On the other hand, continuing to forage during inclement weather could put animals at risk. One major risk faced by many species in arid and semi-arid environments is thermal stress (Smit *et al*. 2016), which can lead to hyperthermia-induced fatigue (Nybo, Rasmussen & Sawka 2014) to the point of being lethal (Saunders, Mawson & Dawson 2011). Moreover, negative correlations with body mass (Costa, Croxall & Duck 1989) and physiological stress (Gesquiere *et al*. 2008) suggest that compensatory foraging effort incurs energetic costs. For example, southern pied babblers (*Turdoides bicolor*) continued foraging despite increasing temperatures, resulting in body mass loss, likely due to reduced foraging efficiency as more breaks for thermoregulation were needed (du Plessis *et al*. 2012). Here, we substantially extend the time window across which we measure the foraging *vs*. thermal homeostasis trade-off (> 2 years) relative to previous studies, and our long-term database (> 5 years) confirms that periods of resource scarcity correspond to body weight loss.

A key remaining question is what exactly drives body weight loss. It could be that, as in the southern pied babblers (du Plessis *et al*. 2012), vulturine guineafowl can sustain foraging activity but pay a thermoregulatory ‘foraging efficiency cost’, meaning that they are less effective during the time that they have available. It is also possible that food becomes so scarce that any (physiological permissible) increase in their foraging time would not be enough to make up for the deficit. Ultimately, these foraging strategies (and levels of thermal risk-taking) were shaped by selective pressures of an evolutionary past that might differ from future climate change contexts (Sih 2024), raising the question whether continuing to forage under high temperatures and low resource availability is still a strategy that pays off in terms of fitness (*sensu* ‘evolutionary mismatch’) when dry seasons become protracted (e.g., during climate-change driven extended droughts). Concerningly, from a conservation perspective, chronic exposure to high temperatures that lead to sublethal fitness costs, including weight loss, can have major downstream consequences for population viability, such as if it contributes to a reduction in breeding. Female birds require good body condition to overcome the burden of reproduction (Moreno 1989; Siikamäki 1998) and birds in our population did not breed (*unpublished data*) during the declared drought that spanned 2021 to 2023 (UNDRR, 2024). Similar impacts may also be expressed in other life-history events that require good body condition, such as migration (Cooper, Sherry & Marra 2015), natal dispersal (Azpillaga, Real & HernándezLMatías 2018) or mate acquisition (Chastel, Weimerskirch & Jouventin 1995). These sublethal fitness costs increasingly threaten birds exposed to changing climates and could lead to widespread population decline and species disappearance (Conradie *et al*. 2019).

We found sex-differences in responses to resource scarcity, with males ramping up foraging more than females and experiencing less weight loss. Sex differences in behaviour or physiology could be critically important if they make one sex more vulnerable to climate change than the other. Foraging strategies under inclement conditions could be driven by sexual dimorphism in body size. For instance, male great bustards (*Otis tarda*) struggle to dissipate heat during hot summer months, resulting in less intensive foraging, relative to smaller-bodied females (Alonso, Salgado & Palacín 2016). In contrast, in Japanese cormorants (*Phalacrocorax capillatus*), larger-bodied males have an advantage in years of food scarcity as they can dive deeper and exploit harder-to-reach foraging areas (Ishikawa & Watanuki 2002). In vulturine guineafowl, males are heavier than females (Table 4), but—counter to what would be expected based on thermal dissipation capacity (Watson & Kerr 2025)—males ramp up foraging more than females. Note here, however, that males are heavier by ∼15%, a difference that might be less consequential than in great bustard where males weigh almost twice as much as females (Alonso, Salgado & Palacín 2016). We instead propose that the sex-modulated effects in foraging activity and weight loss in vulturine guineafowl are more likely driven by differences in life-history strategies. Specifically, females must forage more intensively than males during wet seasons to gain weight needed for egg production and incubation (Nyaguthii *et al*. 2025). During this time, males perform elaborate courtship displays (Guan *et al*. 2022), mate guard and spend considerable time vigilant and/or competing with other males (Nyaguthii *et al*. 2025). In support of this, we found that females spend more time foraging than males during periods of high resource availability (Fig. 2a, see also: Lewis *et al*. 2015). As resource scarcity increases, however, males ‘overtake’ females in the proportion of time spent feeding (Fig. 2a) and showed less dramatic body weight loss (5%) relative to females (9%; Fig. 2b). These differences in slope could again be related to different life-history pressures and social factors. Unlike females who put on weight rapidly in a relatively short time window before laying eggs, males often attempt to maintain body condition prior to the breeding season, which would improve their competitive ability against other males and their access to females (Ball & Ketterson 2008; Vezina & Salvante 2010). Vulturine guineafowl males are also dominant over females, likely increasing their access to resources (Dehnen *et al*. 2022; Papageorgiou, Nyaguthii & Farine 2024). As periods of resource scarcity become protracted, females may have a larger deficit to make up for when favourable conditions return. This would be in line with the pause in breeding in our study population, where females may not have had the physical condition— or the time to build condition back up—to breed even during temporary reprieves in resource scarcity (Fig. S1). Predicting the impact of climate change on animal populations will need to account for how trade-offs faced during harsh conditions carry over to sex-specific life history stages.

Our study adds to an emerging set of studies that are deploying new technology (accelerometers and other sensors) to quantify how wildlife is responding to climate change (Hammond, Palme & Lacey 2018; Noonan *et al*. 2018; Tatler *et al*. 2021). While many papers across a wide array of animal systems have been published on the classification of behaviour (including foraging activity) from accelerometery signals (see: Brown et al., 2013 for review), only a handful of these have used this approach to test how foraging activity budgets shift across environmental gradients (Hernández-Pliego *et al*. 2017; Chimienti *et al*. 2021; Ullmann *et al*. 2023). Together with our study, these demonstrate the value of applying accelerometery data to gain a better understanding of how animals manage foraging *vs* risk trade-offs. These may give crucial insights into the expression of behavioural flexibility and its potential evolutionary mismatch in the context of a future with extended droughts.

## Acknowledgements

We thank the Vulturine Guineafowl Research Programme field team: Brendah Nyaguthii, Wismer Cherono, Monicah Wambui, John Wanjala, Mary Waithira Ngugi, Eunitah Makokha, Anne Namaemba and Micah Wangai, for deploying tags, downloading ACC and temperature data. We thank Christina Hansen-Wheat, Daniel Zuñiga, Edel Odhiambo and James Klarevas-Irby for their contributions to the internal body temperature data, and David Rozen-Rechels for establishing the operative temperature and NDVI data collection and for feedback on this manuscript. We thank Mpala Research Centre for logistical support and allowing us to work on the conservancy. We thank National Museums of Kenya (NMK), the National Commission for Science, Technology and Innovation (NACOSTI) and the Wildlife Research and Training Institute (WRTI) for approval of our work. This research was funded by the ERC (grant no. 850859 awarded to DRF) and by an Eccellenza Professorship Grant of the SNSF (Grant Number PCEFP3_187058 awarded to DRF). CC received additional support from a UZH POSTDOC grant (no. K-74312-02-01).

## Author Contributions

CC and DRF conceptualised the study. KS led the accelerometer and temperature data collection in the field. CC ran the statistical analyses and wrote the first manuscript draft, with input of DRF on both. All authors read and approved the final version of the manuscript.

