## Supplementary material for "Resource scarcity increases foraging activity despite thermal risk in an arid-adapted bird"

**Supporting information**

Data collection

*Section S1: Accelerometer sampling regime*

Daily feeding activity was calculated using tri-axial accelerometer data recorded on solar-powered e-obs tags between 06:20 and 11:30 for n=49 birds (Fig. S1). Tags were programmed to switch to lower sampling rates when battery levels reach a critical minimum threshold, meaning that some days did not collect continuous second-by-second data between 06:20 and 11:30 (310 minutes). Moreover, accelerometer data collection is interrupted while the GPS is finding a satellite connection and taking a fix every 5 minutes (e-Obs system manual). Thus, we included mornings if they had at minimum 155 minutes of accelerometer data (median: 288 minutes; range: 156 – 305 minutes). Further, a feeding supplementation experiment (Ogino 2025) took place between 2022-06-03 and 2022-07-12 (Fig. S1) that likely affected the feeding behaviour of the birds, hence we excluded data from this period from our analyses.

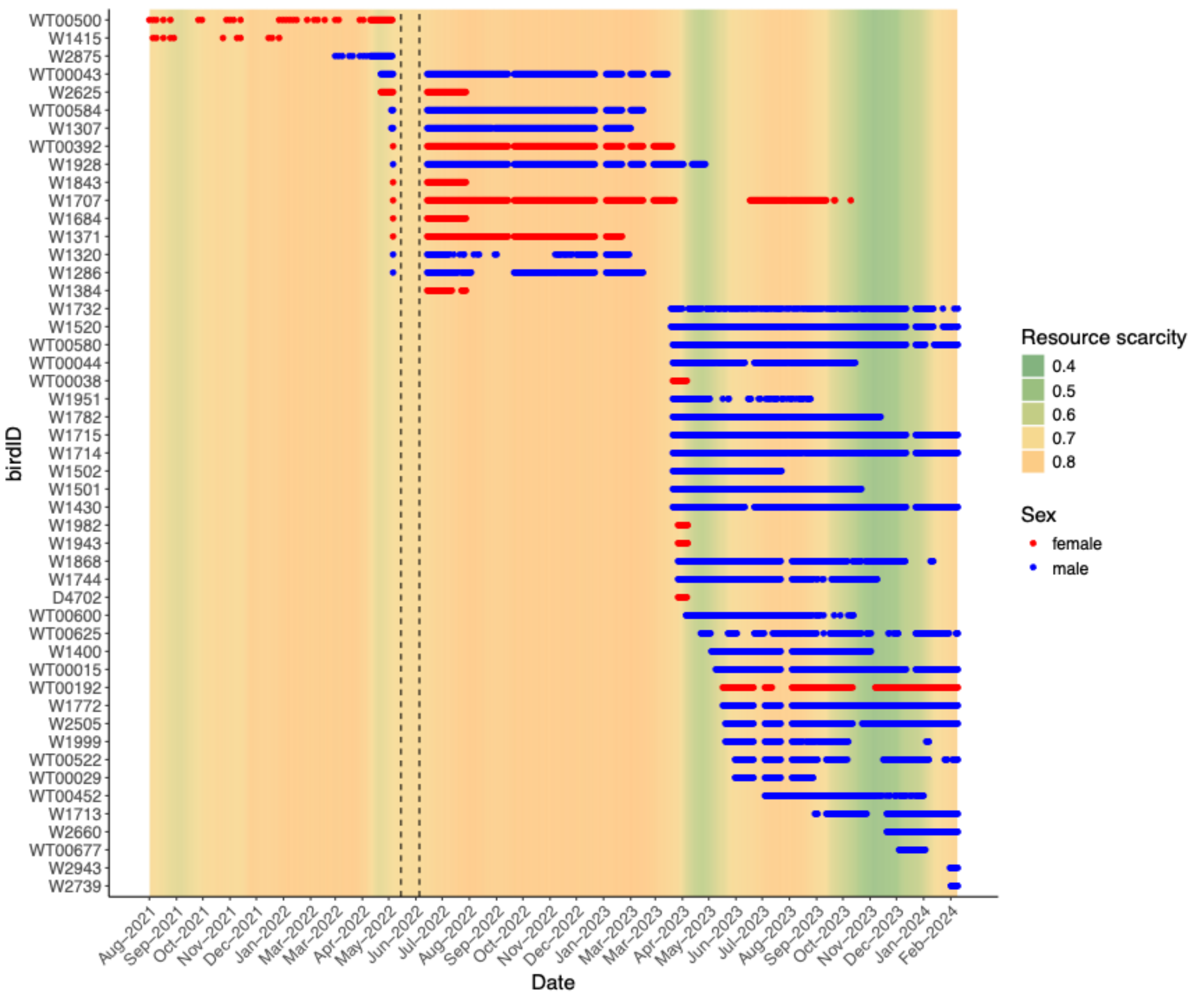


**Figure S1**. Temporal distribution of data for birds fitted with biologging tags collecting accelerometer data (sex denoted by blue = male; red = female) and the coinciding environmental conditions in the background (colour scheme from green to orange indicate increasing resource scarcity (1-NDVI). Black dashed vertical lines denote the period where the feeding experiment took place (Ogino, 2025) which was removed from the accelerometer dataset for analyses. Note that early 2021 through to mid 2023 was a severe drought (UNDRR, 2024). Each dot is a day of data with gaps representing missing data days, due to low tag battery or the tags being pre-emptively reprogrammed to pause accelerometer-data collection to avoid batteries depleting.

*Section S2: ECG logger data inspection*

From inspection of the internal temperature logger data of n=7 birds implanted with ECG loggers (Brandl et al., 2025), one bird was identified as having outlier temperature readings (mean±SE = 39.4±1ºC) relative to the others that had more similar ranges (mean ± SE = 40.5±0.6ºC; Fig. S2a). The latter values also corresponded more closely to the ranges reported for other fowl: broiler chickens *Gallus gallus domesticus* between 41.2 - 42.2ºC (Hamrita & Conway 2017); free-ranging helmeted guineafowl *Numida meleagris coronata* 40.5-44.5ºC (Van Niekerk, Megía-Palma & Forcina 2022); farmed helmeted guineafowl *Numida meleagris* 39.9-43.8ºC (Ayo *et al.* 2022). As we could not determine whether this 'outlier bird' was caused by measurement error or true biological differences, we excluded it (but Fig. S2b for result with 'outlier bird'). Our resulting sample size was therefore six birds for a total of 630 bird days (median: 108, range: 29-155 days of logging per bird).

*Section S3: NDVI calculation*

We use Normalized Difference Vegetation Index (NDVI) data (resolution: 10 by 10 meter cells) calculated from European Space Agency Copernicus Sentinel-2 satellite images to assess change in vegetation quality (Pettorelli *et al.* 2005). In brief, we downloaded the imagery tile covering the region and computed the NDVI for each cell. We then sub-set the cells to a polygon that corresponds to the population footprint between 2018 and 2024. Next, we identified the cells with cloud cover, removed them, and calculated the mean NDVI across remaining cells within the polygon. We removed days with outlier values (NDVI <=0.09), which were caused by excessive cloud cover (confirmed visually from the images). NDVI data were collected every 5$^{th}$ day, and we smoothed these values using a Gaussian smoother (which increases the weighting of the current day and decreases the weighting of days further away in the smoothing window) before interpolating the values to obtain daily NDVI values (to correspond to the sampling regime of the biologging data).

Data analysis

*Section S4: Identifying a 'thermal threshold' in vulturine guineafowl*

To identify a 'thermal threshold'—the operative temperature at which the internal body temperature reaches a maximum—we matched the mean hourly internal body temperature (from ECG loggers, see *Section S2*) for each bird to the corresponding mean hourly external operative temperature measured by external temperature loggers. We then fit a linear mixed model with internal body temperature as the response and external operative temperature as a second order polynomial term, reflecting a scenario where internal temperature can increase with operative temperature until at a certain temperature limit, whereupon it stabilises or drops again due to physiological or behavioural adjustments made by the animal to avoid hyperthermia. We included for sex as an interactive term, as sex differences in bird body temperatures have been reported (see: Prinzinger, Preßmar & Schleucher 1991, for review) and controlled for bird ID and date as random intercepts. We scaled external operative temperature from 0-1 to allow for interpretation alongside sex. We arrived at the polynomial model using backward model selection based on a likelihood ratio test and we dropped sex as it did not significantly predict differences in internal body temperature (Table S1). To calculate the maximum point of the curves, we extracted the coefficients of the first-order term (a; linear coefficient) and the second-order term (b; polynomial coefficient) from the model and used the vertex formula for a parabola to find the x-coordinate:

$$\textit{operative temperature at maximum body temperature} = -a / (2 * b)$$

Further, we took the intercept of the model (*i*) and used the following formula to find the associated y-coordinate:

$$maximum\ internal\ body\ temperature = i + a * max.\ operative\ temp. + b * (max.\ operative\ temp.)^2$$

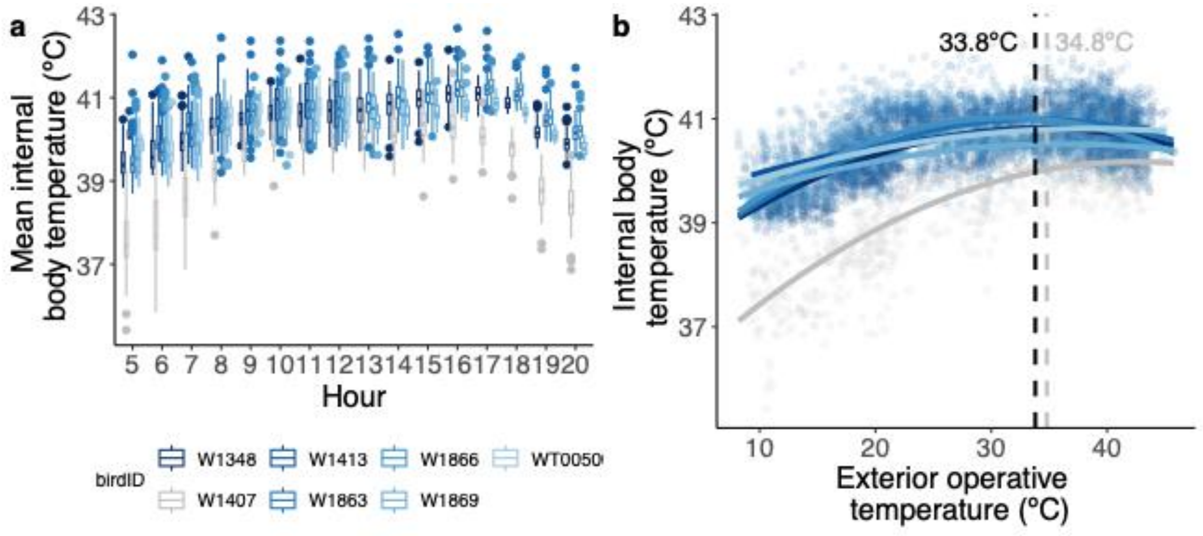


**Figure S2**. (**a**) Internal body temperature per hour for n=7 vulturine guineafowl between the hours of 05:00 and 20:00 EAT. W1407 (in grey) was removed from the analysis due to anomalous temperature readings relative to the other n=6 birds and fowl body temperatures reported in the wider literature. (**b**) Calculation of maximum values (indicated by dashed vertical lines) the exterior operative temperature reached before internal body temperature decreased again when including the outlier bird (W1407) in grey and without the outlier bird in black.
